# Modulation of error sensitivity during motor learning across time, space and environment variability

**DOI:** 10.1101/2025.05.30.656374

**Authors:** Dmitrii I. Todorov, Maelys Moulin, Ethan Buch, Romain Quentin

**Affiliations:** EDUWELL Team, Lyon Neuroscience Research Center (CRNL), INSERM U1028, CNRS UMR5292, Université Claude Bernard Lyon 1; Sorbonne Université, CNRS, Inserm, Laboratoire d’Imagerie Biomédicale, LIB; IDH Team, LIRMM, CNRS, University of Montpellier; EuroMov Digital Health in Motion, University of Montpellier, IMT Mines Alès; Human Cortical Physiology and Neurorehabilitation Section, NINDS, NIH

**Keywords:** Motor adaptation, Reaching movements, Visuomotor rotation, Error sensitivity, Adaptation rate, Generalization, error-based adaptation, Motor learning, Movement variability

## Abstract

We never experience the exact same situation twice. In our dynamic and constantly changing environment, we continuously need to adapt our behavior, either due to external (e.g., a change of wind) or internal factors (e.g., muscle noise). Motor adaptation is the process of recalibration of movements in response to such perturbation. It activates when an agent needs to change their movement in response to a perceived error. It has been proposed that in order to perform motor adaptation, the brain continuously updates an error sensitivity signal that controls how much is learned from a past error. Such error sensitivity reflects the speed of learning, and understanding its dynamics is crucial to promoting efficient learning. In our experiment, healthy participants performed a visuomotor adaptation task requiring reaching multiple targets using a joystick. They experienced two periods of stable perturbations and two periods of random perturbations. In agreement with past research, we found that error sensitivity is lower in unstable environments. Moreover, error sensitivity is higher in a stable environment without perturbation than in a stable environment with perturbations. We observed a continuous increase of the error sensitivity within a stable perturbation environment. Finally, continuous spatial (across target locations) and temporal (across trials with same target) generalization of learning is present not only in stable but also in the random environment.

## Introduction

Motor learning corresponds to the improvement of motor skills through practice. It is an essential part of human behavior. Motor adaptation is a particular type of motor learning where an agent adjusts their movements through trial and error and in response to a change in the environment (Malone, Vasudevan, and Bastian 2011). In humans performing reaching movements, the motor adaptation process is largely driven by the perceived error (Mazzoni and Krakauer 2006; Morehead and Orban de Xivry 2021). The degree to which such errors are used to update the next movement is determined by so-called *error sensitivity* (ES) (Herzfeld et al. 2014) –– that is, an error-and-time-dependent adaptation rate. While many studies have initially assumed that the error sensitivity is constant (Donchin, Francis, and Shadmehr 2003; Jordan and Rumelhart 1995; Kawato, Furukawa, and Suzuki 1987; Thoroughman and Shadmehr 2000), more recent studies have investigated its dynamics and propose computational models that describe how error sensitivity is updated based on past behavior. For example, it was posited that the update of error sensitivity depends on the sign consistency between two consecutive errors, with error-sensitivity increasing when two two consecutive errors have the same sign) (Herzfeld et al. 2014). Another model suggested that error sensitivity is dynamically modulated, depending on the mean and standard deviation across a temporal window of previous errors (Tan, Jenkinson, and Brown 2014).

A crucial phenomenon in motor adaptation is *generalization*. When adapting to a new environment, we rarely start from zero. Generalization is the transfer of knowledge from one context to another and it is a fundamental part of the motor adaptation process (Krakauer et al. 2006). At least two types of generalization exist: Across space, in which error information from a movement to one target influence the next movement to a different target, and across time, in which error information from an earlier trial influence a later movement, even if other trials occurred in between.. Prior studies of spatial generalization have investigated this phenomenon from many different angles but always assuming static learning rate (Krakauer et al. 2006; Tanaka and Sejnowski 2014). The generalization of learning between two spatially distinct targets is modulated by the distance between them (Donchin et al. 2003). The closer the two targets are, the greater the extent of adaptation generalization. Decay of generalization as a function of distance between targets is not solely due to muscle patterns being different for different targets for force field (Berniker et al. 2014) and digitizing tablet (Brayanov, Press, and Smith 2012). Very few studies have investigated temporal generalization (Donchin et al. 2003). In this study, we investigated how error sensitivity dynamically changes depending on context stability, and examined its generalization across time and space.

Only few studies investigated such trial-resolved dynamics of error sensitivity and only a handful of perturbation schedules were explored. In our study, twenty participants performed a visuomotor adaptation task (Fig. 1) in which they had to reach one out of four possible targets by controlling a cursor with a joystick at each trial. We specifically studied how error sensitivity evolves within environmental conditions with predictable (“stable”) perturbation and with completely unpredictable (“random”) perturbations. Error sensitivity was larger in stable than in random environment but only when including zero-perturbation trials (Fig. 2C and 2D). Interestingly, error sensitivity in a stable environment was the highest right before the perturbation was introduced, dropped at the onset of a new perturbation (Fig. 2C) and grew during the perturbation or washout, similarly to (Herzfeld et al. 2014) (Fig. 3). Such an increase of error sensitivity during a stable environment survives even when the previous error size is controlled for. Our results also demonstrate that participants generalize their learning from errors across space and time with a monotonous generalization curve, both in stable and in random environments (Fig. 4). We found that, in our data, error sensitivity values are not fully consistent with two well-known models: error-sign-consistency model (Herzfeld et al. 2014) and error sensitivity being ratio of squared mean and variance inside a temporal window of 20 trials (Tan et al. 2014).

**Figure 1.**
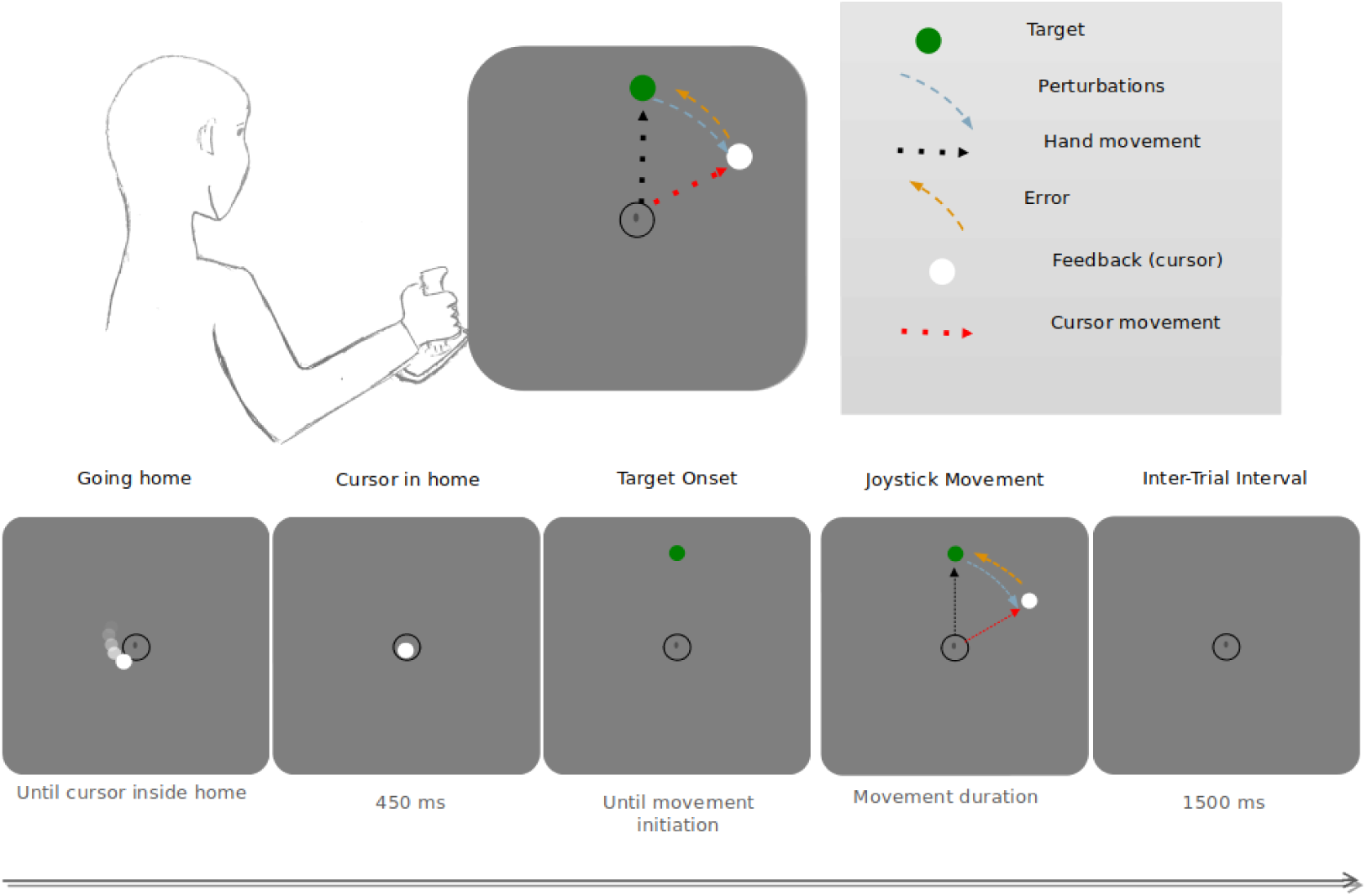
Experimental design. Participants controlled a cursor with a joystick to reach targets appearing on the screen. On some trials, visual feedback was perturbed by a visuomotor rotation of 30 degrees clockwise or counterclockwise.

**Figure 2.**
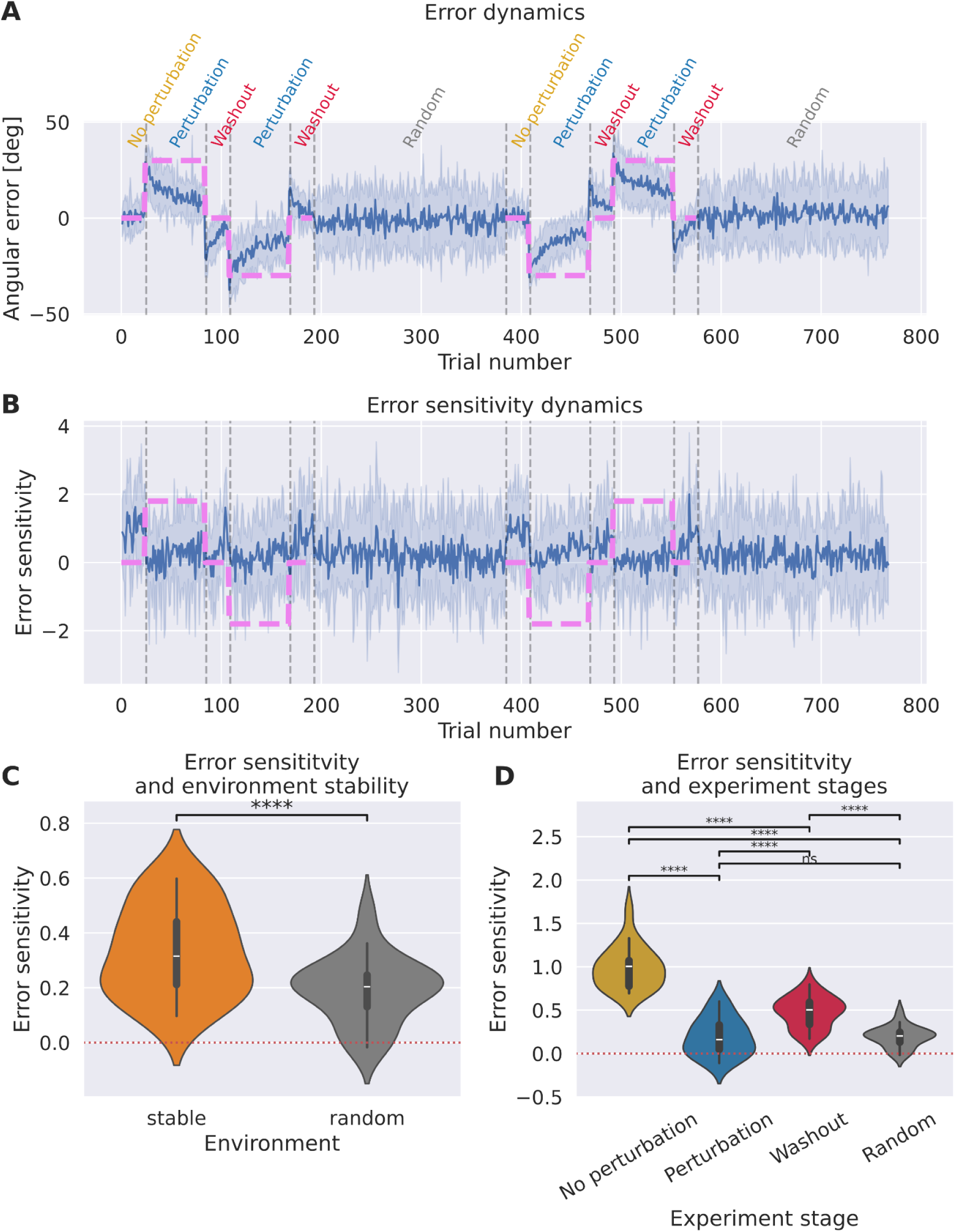
**A**. Error dynamics. **B.** Error-sensitivity dynamics. Both **A** and **B** for each trial show a mean across participants, shading represents standard deviation, purple line represents perturbation, text on top of **A** denotes experimental conditions. **C.** Within-participant average error sensitivity in stable and random environments. Each violin plot is made using 20 points corresponding to values for the respective condition, averaged within each of the participants). **D** same as **C** but with a more fine-grained stable environment, colors are consistent with the text colors in panel A. On both plots **C** and **D,** all values are significantly positive. The p-values codes are after Holm-Bonferroni correction applied.

**Figure 3.**
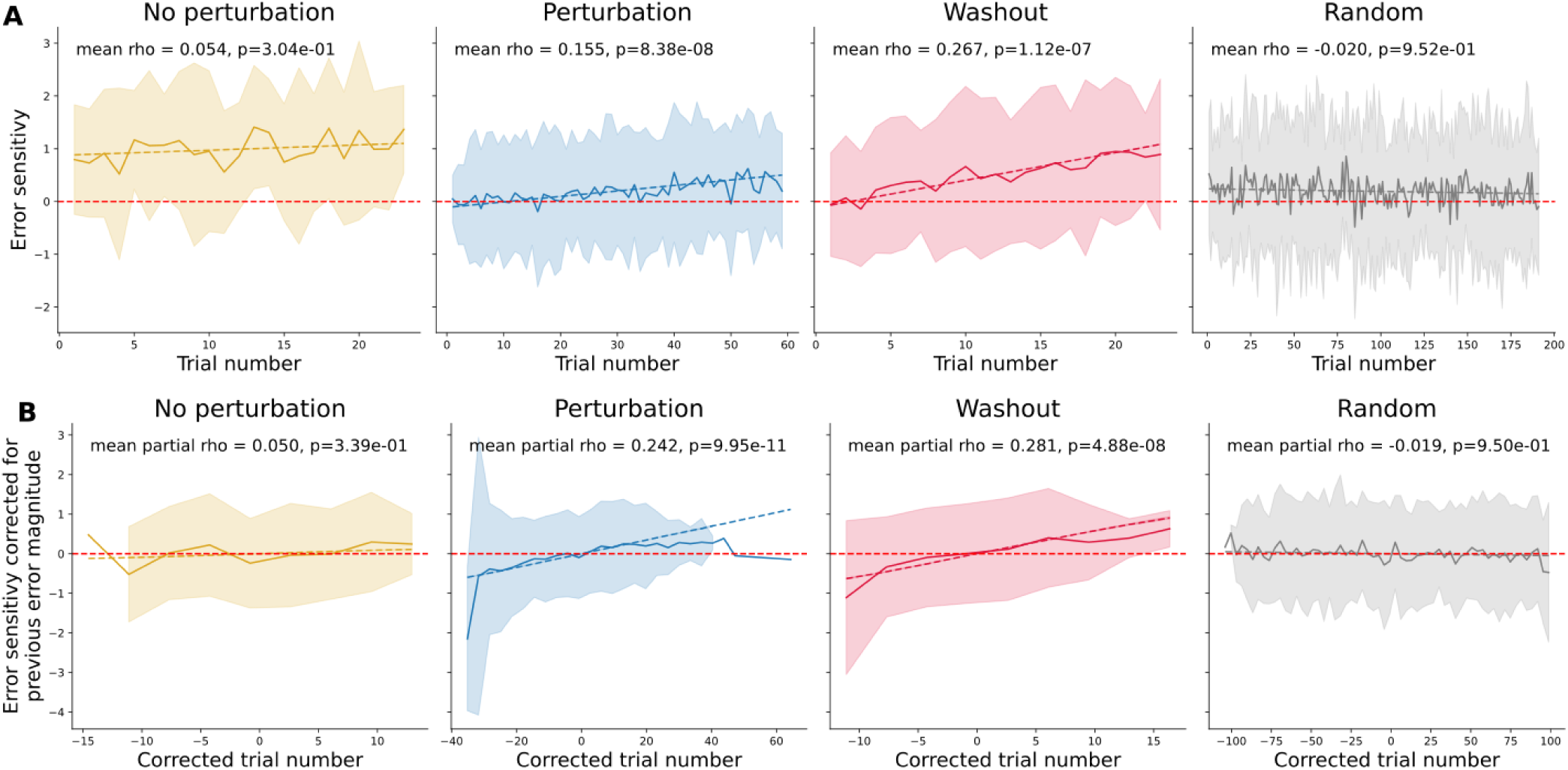
Error sensitivity dynamics in different environments. **A.** Correlations between error sensitivity and trial number in each environment, B: Partial correlations between error sensitivity and trial number, controlled for the absolute error magnitude on the same trial. The corrected trial number corresponds to the residuals of the correlation between trial number and absolute error size and the corrected error sensitivity corresponds to the residuals of the correlation between error sensitivity and absolute error size. Solid lines represent the mean over participants and conditions and shadings represent standard deviations. No perturbation, Washout and Random environments have two sets of data representing two separate appearances of the condition for each subject, Perturbation has four such sets (see Fig. 2A). Dashed lines are linear fits. The mean spearman rho across participants and the significance of the one-sample t-test against zero is annotated for each panel. These results show that error sensitivity grows during perturbation and washout periods.

**Figure 4.**
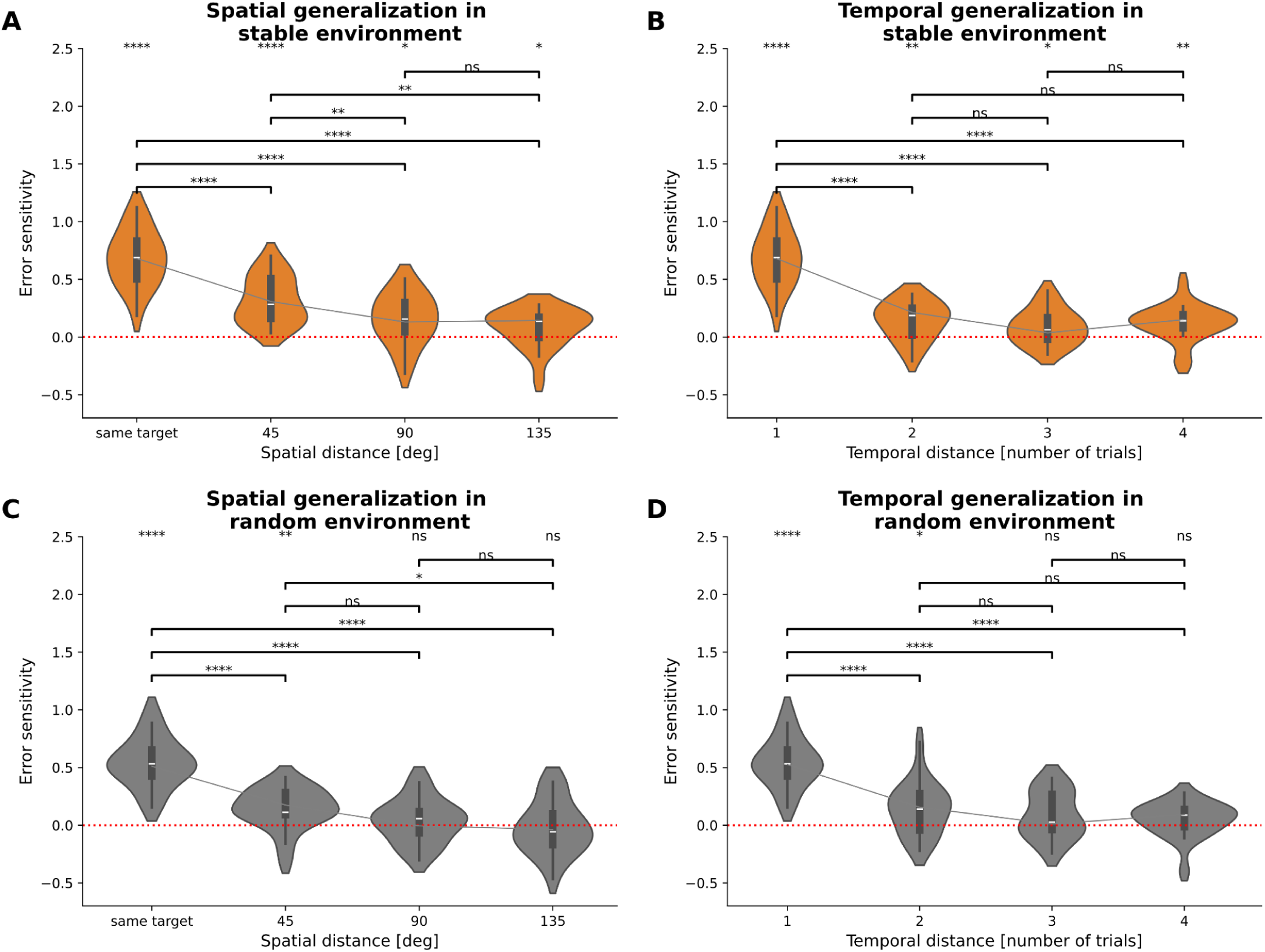
Generalization patterns. **A** and **C**. Spatial generalization in stable and random environments. Error sensitivities are measured separately depending on the distance between targets in two consecutive trials. **B** and **D**. Temporal generalization in stable and random environments. Error sensitivities are measured as the correction of either the preceding error (n-1) or the error at n-2, n-3 or n-4 trials. Error sensitivities are averaged within participants. Each violin plot represents data from 20 participants. Solid line shows a polynomial fit to means. The top line of significance codes are for comparison of error sensitivity with zero. All significance values were computed using Holm-Bonferroni correction. Note that for the spatial distance, the average error sensitivity within distances are computed using different numbers of trials because targets were uniformly distributed and fewer trials have larger target jumps between consecutive trials.

Overall these results improve our understanding of dynamic motor adaptation, which in turn could help developing improved motor rehabilitation protocols for conditions such as stroke in the future. (Krakauer 2006; Krakauer and Carmichael 2022).

## Methods

### Participants and experimental sessions

Twenty healthy and right-handed volunteers (15 women, 5 men, mean age of 25.9 +/− 4.1 years) participated in the study after providing their written informed consent to participate in the project, which was approved by the Combined Neuroscience Institutional Review Board of the National Institute of Health (NIH). They all had normal physical and neurological examinations and normal or corrected-to-normal vision. This study was not preregistered.

### Visuomotor adaptation task

Visual stimuli were displayed using pygame Python 3 package (https://www.pygame.org). The visual stimuli were back-projected on a translucent screen in front of the participants. The screen was positioned at a distance of 80 cm from both eyes of the participants. Each trial started with a black circle (called the home, radius = 1.5 cm) at the center of the screen (Fig. 1). Participants were instructed to position a white cursor (filled circle, radius = 0.6 cm) inside the home position using a joystick fiber optic response pad (Current Designs, Inc., USA) with their right hand. After maintaining the cursor position inside the home position perimeter for 464 ms, a green target (filled circle, radius = 0.9 cm) appeared at one out of 4 possible positions (22.5°, 67.5°, 112.5° and 157.5 on a unit circle centered on the home position with a radius of 20 cm), while the white cursor simultaneously disappeared. Participants then used the joystick to move the cursor towards the target position. The (unperturbed) cursor position coordinates were a simple linear transformation of the two joystick outputs (horizontal and vertical angles) provided by the joystick driver. The visual feedback of the cursor was absent during the joystick movement, only reappearing as a static endpoint feedback once the home-to-cursor distance reached 20 cm (i.e., passed through the invisible target array circle), regardless of whether the subject hit or missed the target. The movement lasted on average 750+/−190 ms. Participants used this endpoint feedback, which was presented for a total duration of 232 ms, to perceive their error (i.e., the target-to-end-point cursor feedback distance). After an inter-trial interval of 1500 ms, participants returned the cursor to the home position with the joystick. The cursor had to remain again within the home position perimeter for 464 ms in order to trigger the next trial target presentation. The entire trial lasted on average 3.729 +/− 0.332 s. The task was composed of four blocks of 192 trials each (768 trials in total) and lasted on average 50.3+/−5.07 minutes. A stable environment block (Fig. 1) consisted of 24 trials without perturbation, followed by 60 trials with a +30° (for half of the participants) or –30° (for another half) visual perturbation, followed by 24 trials without perturbation, followed by 60 trials with a –30° (or +30° if the first perturbation was –30°) and followed by 24 trials without perturbation. Participants were pseudo-randomly assigned to start with a +30° or –30° perturbation. In the second and the fourth blocks, the perturbation was applied pseudo-randomly over the 192 trials. More precisely, errors presented during these blocks were randomly permuted replays of the errors experienced by the participant in the previous stable block. Thus, end-point cursor feedback presented on each trial during the random perturbation block had no fixed relationship to the actual movement performed (such trials are often called “error clamp” trials (Smith, Ghazizadeh, and Shadmehr 2006)). This design approach ensured that the overall distribution of errors was controlled and kept consistent between the stable and random environments.

### Data analysis

Half of participants started with a +30° perturbation and another half started with a –30° perturbation. To simplify group analysis, we inverted the signs of errors, reach angles and target angles for participants who started with the –30° perturbation and further conducted the analyses only with these values.

We define error sensitivity at the beginning of trial *n* using the following formula, derived from (Herzfeld et al. 2014).

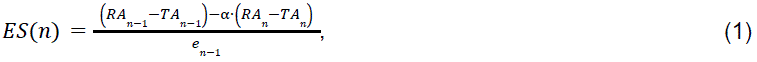

where *RAn*−1 and *RAn* are the angles between the endpoints of the subject movement and the origin *before visuomotor perturbation is applied* at trials *n* – 1 and *n*, and *TAn*−1 and *TAn* are the target angles at these trials (target location is never perturbed by the visuomotor perturbation but the target locations generally change from trial to trial), *en*−1 is the perceived visual error at the feedback onset of trial n-1 and α is the retention factor. We emphasize that here, *ES*(*n*) represents the value of error sensitivity at the start of the trial, though it is determined post-hoc by analyzing the reach angle at the trial’s end. Formula (1) corresponds to formula (s5) in (Herzfeld et al. 2014). We used the value α = 0. 924 following (Avraham, Keizman, and Shmuelof 2020).

All statistical analyses were performed in python using pingouin package v0.5.5 (Vallat 2018).

#### Hit counts

A trial is considered a “hit” when, at the end of the trial, the displayed cursor circle intersects with the target circle. Participants hit the target on average in 15.7%, ± 2.9% of all trials. Mean hit proportions across environments are identical because errors during random blocks were drawn from the preceding stable block.

We remove trials for which previous trials were hit trials from consideration because there is no error to correct in the next movement. We also remove trials where error sensitivity exceeds 5 times the mean standard deviations within participants (corresponding to 0.85% of remaining trials after excluding trials with a preceding hit.

#### Data Availability Statement

Data (subject to participant consent) used to generate the findings of this study are freely available at OSF (Todorov 2025): https://doi.org/10.17605/OSF.IO/6J8SM. Custom code generated during this study is available at <u>github.com</u>.

#### Use of Artificial Intelligence (large language models-enabled tools)

During the preparation of this manuscript, generative artificial intelligence (AI) models were utilized as productivity tools. The authors take full responsibility for the accuracy, integrity, and originality of the final content. The AI models used include Google AI Studio (Gemini 2.0 Pro, Gemini 2.5 Pro, Gemini Flash 2.5), OpenAI’s ChatGPT-4o, Google NotebookLM, Elicit, Scite, Perplexity, and integrated coding assistance models within Visual Studio Code (VSCode Copilot). No raw data was analyzed by AI tools directly.

The application of these tools can be categorized as follows:

● Code Development and Debugging: AI was used to accelerate the development of Python scripts for debugging and figure generation. All AI-generated code was thoroughly reviewed and validated manually by the authors.
● Manuscript Formatting: AI assisted in checking grammar and syntax and in rephrasing sentences where required.
● Literature Discovery: AI web search tools (specifically Elicit, Scite and Perplexity) were used to check for the existence of relevant literature. These tools use web search to avoid hallucinating non-existing papers’ names.
● Document Analysis: AI models capable of “smart search” (such as Google NotebookLM) were used to locate and answer precise questions about the content of specific research papers loaded into the tool. All information retrieved in this manner was subsequently verified by the authors against the original source documents.

## Results

### Error Sensitivity is higher when no *external* perturbation is present

Error sensitivity is predominantly positive in all environments (Fig. 2D, one-sample t-tests against zero, all p<0.05, during no perturbation condition: error sensitivity = 0.99 ± 0.24, 95% CI [0.9, inf], t(19)=18.29, p=3.22e-13, d=4.09; during perturbation: error sensitivity = 0.19 ± 0.22, 95% CI [0.11, inf], t(19)=3.99, p=3.92e-04, d=0.89; during washout condition: error sensitivity = 0.48 ± 0.18, 95% CI [0.41, inf], t(19)=12.18, p=3.01e-10, d=2.72; during random condition: error sensitivity = 0.19 ± 0.12, 95% CI [0.15, inf], t(19)=7.19, p=7.87e-07, d=1.61). All p-values are Holm-Bonferroni corrected for multiple comparisons. It demonstrates that participants compensate for their errors even in the random environment. Negative error sensitivity values are occasionally observed at the trial level, which can be attributed to movement noise when participants use a joystick to control a cursor.

Error sensitivity is significantly higher in stable (mean=0.33±0.16) than random (mean=0.19±0.12) environment (two sample t-test, mean difference = –0.14, 95% CI [[-0.19 –0.08]], t(19)=-5.19, p=5.24e-05, d=0.96) (Fig. 2C).

To compare error sensitivity in sub-conditions, one way repeated measures ANOVA revealed a significant effect of condition on error sensitivity (Greenhouse-Geisser corrected F(1.98, 37.63) = 85.70, p < .001, generalized eta-squared = 0.75), see also Table S2. We then conducted post-hoc pairwise comparisons using t-tests with Holm-Bonferroni corrections. Error sensitivity sharply increases when no perturbation is present. Specifically, as visible on Fig. 2B, error sensitivity is significantly higher during the no perturbation period (mean=0.99±0.24) compared to periods with stable perturbations (mean=0.19±0.22) (mean difference = 0.80, 95% CI [0.67, inf], t(19)=10.55, p=1.33e-08, d=3.49). Error sensitivity is also higher during washout periods (mean=0.48±0.18) compared to periods with stable perturbations (mean difference = 0.28, 95% CI [0.19, inf], t(19)=5.38, p=2.06e-04, d=1.45). Surprisingly, error sensitivity during stable perturbations is not significantly different from the error sensitivity in the random condition (t=0.05, p=9.59e-01).

These results highlight the influence of environmental stability on the learning process.

### Error sensitivity grows within stable environment

Our results demonstrate that error sensitivity grows with time during perturbation and washout periods of stable environment (one-sample t-test against zero on the distribution of spearman rho across participants, for the perturbation condition: rho=0.15 ± 0.08, 95% CI [0.12, inf], t(19)=8.77, p=8.38e-08, d=1.96, for the washout condition: rho=0.27 ± 0.14, 95% CI [0.21, inf], t(19)=8.44, p=1.12e-07, d=1.89). To account for the potential influence of error magnitude on these correlations, partial correlations conditioned on the absolute error size of the previous trial error were also examined and results were similar (for the perturbation condition: rho=0.24 ± 0.08, 95% CI [0.21, inf], t(19)=13.22, p=9.95e-11, d=2.96, for the washout condition: rho=0.28 ± 0.14, 95% CI [0.23, inf], t(19)=8.91, p=4.89e-08, d=1.99). All p-values in this paragraph are Holm-Bonferroni corrected.

The perturbation condition exhibited a stronger positive correlation compared to the random condition both for classic correlations (paired two-sample t-test comparing rho values: mean difference = 0.17, 95% CI [0.14, inf], t(19)=9.30, p=1.01e-07, d=2.637) and partial correlations (mean difference = 0.26, 95% CI [0.23, inf], t(19)=12.76, p=5.49e-10, d=3.85). In the washout condition the correlation and partial correlation were larger than in no perturbation condition (for regular correlation mean difference = 0.21, 95% CI [0.11, inf], t(19)=3.53, p=1.12e-02, d=1.12, for partial correlation mean difference = 0.23, 95% CI [0.14, inf], t(19)=4.34, p=1.76e-03, d=1.23). Unsurprisingly rho values in washout are larger than in the random condition as well (for regular correlation mean difference = 0.29, 95% CI [0.23, inf], t(19)=8.19, p=6.53e-07, d=2.70, for partial correlation mean difference = 0.30, 95% CI [0.24, inf], t(19)=8.63, p=2.95e-07, d=2.84). All p-values in this paragraph are Holm-Bonferroni corrected.

### Generalization across time and space in stable and random environments

Characterizing how error sensitivity is influenced by the temporal distance between trials and the spatial distance between targets provides insights into the nature of motor memory and adaptation across different conditions. Spatial generalization is a phenomenon of transfer adaptation across nearby movements and more specifically, across nearby reaching targets. To conduct spatial distance analysis, we grouped trials by the distance between the current and the previous trial target (see Figure S8). Error sensitivity decreases with spatial distance both in stable and random environment (in stable: rho = –0.77 ± 0.36, 95% CI [-inf, –0.63], t(19)=-9.49, p=6.08e-09, d=2.12; in random: –0.77 ± 0.28, 95% CI [-inf, –0.66], t(19)=-12.09, p=1.15e-10, d=2.70), indicating that errors made while reaching for spatially distant targets influence less the learning than errors associated with proximal targets.

Temporal generalization assessed sensitivity not only to the most recent error but also to preceding errors. To conduct temporal generalization analysis, we can define an extension to formula (1): instead of dividing by the most recent error we can divide the error correction by the k-preceding error in the following way

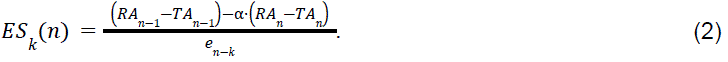

This quantity shows how correction on trial n is modulated by error history of depth k. We observed a continuous decline in error sensitivity with increasing temporal distance both in stable and random environment (in stable: rho = –0.62 ± 0.32, 95% CI [-inf, –0.49], t(19)=-8.56, p=3.01e-08, d=1.91; in random: rho = –0.65 ± 0.30, 95% CI [-inf, –0.54], t(19)=-9.80, p=3.61e-09, d=2.19, indicating that errors occurring in earlier trials contribute less to learning than more recent errors, regardless of the target’s consistency.

Note that there are two ways one can use the quantity in formula (2): either by looking at *all trials* or by restricting to the subset of trials for which the *target at trial n-k was the same as during trial k*. We report above the latter (keeping the same target) because the former shows no evidence of generalization in the random environment (see Fig S1) and for the stable environment the results remain qualitatively the same.

To check whether the environment stability affects the generalization across space, we have computed two-way repeated measures ANOVA. The interaction effect between environment and the distance between the consecutive targets was not statistically significant, F(3, 57) = 0.42, p = 0.686, generalized eta-squared = 0.004, see also Table S3. This suggests a consistent generalization pattern across conditions.

### Error sensitivity grows when recent errors have consistent signs and drops when they don’t

The paper that coined the term “error sensitivity” (Herzfeld et al. 2014) proposed a formula for its trial to trial evolution. Specifically, the formula (3) in (Herzfeld et al. 2014) proposes that between two consecutive trials the error sensitivity changes additively: grows if the previous trial errors had the same sign and decreases if the signs were different (see also Discussion). We can test this in our data directly by checking how our empirically defined quantity from formula (1) changes between consecutive trials in relation to consistency of consecutive error signs. Formally we define the error sensitivity change with the following formula:

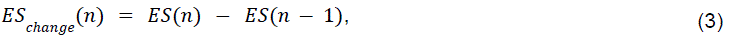

In this analysis, trials for which either previous or the pre-previous trial was a hit trial were removed (otherwise error sensitivity values are too high and error sensitivity change is not informative). In average, 15% ± 3% of trials per participant were removed (in addition to those above 5*std threshold removed for all analyses in the beginning). In the stable environment, when consecutive errors are inconsistent in sign, there is a decrease in error sensitivity, with a mean change = –1.14 ± 0.26, 95% CI [-inf, –1.04], t(19)=-19.29, p=3.06e-14, d=4.31. Conversely, for consistent consecutive errors, error sensitivity increases, evidenced by a mean change 0.31 ± 0.13, 95% CI [0.25, inf], t(19)=10.19, p=1.93e-09, d=2.28.. This corroborates findings of Herzfeld et al (Herzfeld et al. 2014). However, in the random environment, there is no increase of the error sensitivity when consecutive errors are consistent in sign (p=7.20e-01) or decrease of error sensitivity when consecutive errors are inconsistent in sign (p=2.23e-01). These results imply that in the unpredictable conditions of the random environment, the consistency of error feedback does not seem to influence error sensitivity adjustments. Nevertheless, learning persists in this unpredictable setting, as indicated by the above-zero average error sensitivity (Fig. 2D)

It is possible that such results depend on motor variability. Maybe error sensitivity should not be expected to change if the inconsistency of error signs happens only due to motor noise. We checked that by repeating the analysis in this paragraph excluding trials whether either previous or pre-previous errors were smaller than the standard deviation of the error in the no-perturbation stage. Although such trials constitute a large proportion of all trials within participants (mean = 64 %, std = 13 %), after excluding them overall, results remained qualitatively the same (stable inconsistent is negative with p=1.13e-11, and stable consistent is positive with p=1.06e-06, see also Fig. S2).

### Error sensitivity relation to statistics of recent errors

In (Tan et al. 2014) authors suggested that one can use a nonlinear transformation of the perceived errors to compute trial-by-trial adaptation rate. Specifically in (Tan et al. 2014) the history of 20 last trials was considered and the following formula was used:

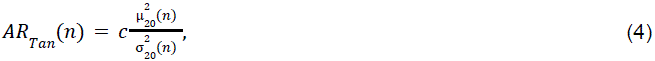

where 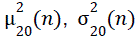 are square of the mean and variance of the observed error in the window of last 20 trials preceding (but not including) trial n, and the coefficient c was fitted using a computational model (note that if one does not ask for the precise value of the coefficient c, the proportionality can be checked without using a computational model). The authors of (Tan et al. 2014) also considered other history lengths and other powers for µ and σ but they were deemed less appropriate.

We tested whether a similar quantity is related to error sensitivity in our data. We compared error sensitivity and *ARTan* by calculating the Spearman correlation between formula (4) with c=1 and formula (1). Spearman correlations were computed separately for stable and random conditions for each participant and spearman rhos distributions were then compared to zero. We did not find statistically significant positive correlations neither in stable, nor in random conditions: we tested the hypothesis of a positive Spearman correlation but contrary to our hypothesis, a one-tailed test for a positive correlation in the stable environment was not significant (p = 1.00e+00). Instead, we found a statistically significant negative correlation (T = –5.44, p < .001). The mean R-squared value for this significant relationship was 0.013. For comparison, no significant positive correlation was observed in the random environment (p = 7.52e-01). Same for negative correlation in the random environment (p = 2.48e-01). See also the scatter plots showing relationship between *ARTan* and error sensitivity in Fig. S6.

One can also modify the expression (4) slightly to produce other kinds of windowed statistics, changing the powers of the mean and standard deviation, similar to how it was done in (Tan et al. 2014). We found, while some of them did show significant correlations in one of the conditions, none of them had significantly positive correlations simultaneously in random and in stable conditions (See Supplementary).

This analysis suggests that the quantity suggested by (Tan et al. 2014) does not seem to influence error sensitivity in our data.

## Discussion

### Summary of results

To investigate the dynamics of error sensitivity during a motor adaptation task, we measured error sensitivity during several conditions, including stable and random environments. We report four main findings. First, error sensitivity is higher during no perturbation than during perturbation, and it grows within a stable environment. Second, error sensitivity is still above zero even in a random environment. It means that participants continue to correct their errors even when nothing can be learned from them. Third, we observed spatial and temporal generalization in both stable and random environments. Fourth, neither the error consistency (Herzfeld et al. 2014) nor windows error statistics-based (Tan et al. 2014) models for error sensitivity dynamics fully agree with our data.

### Dynamics of error sensitivity in stable versus random environments

Error sensitivity during no perturbation conditions is significantly higher than during perturbation and washout conditions (Fig. 2D). This increase, visible at the trial level each time the perturbation disappears (Fig. 2B), could be explained because participants are more eager to compensate for their own motor noise (motor variability present physiologically in the absence of any perturbation), maybe because they are more familiar with it, than with an external perturbation. The error sensitivity also grows within stable perturbation, which one could explain in two ways: either i) towards the end of the perturbation, the internal model adaptation stabilizes and then the same explanation as for no perturbation explains higher error sensitivity or ii) the perturbation onset induces a drop in error sensitivity and then the error sensitivity slowly increases until the next change in stability. The latter is closer to the hypothesis in (Herzfeld et al. 2014).

The washout phase, marked by the removal of perturbations, seems to facilitate a recovery or enhancement in error sensitivity, with a significant positive correlation between trial number and error sensitivity. In contrast, during random conditions, error sensitivity gradually decreases over time, although this decrease is not statistically significant. Such a negative trend suggests that unpredictable conditions may hinder the adaptive process.

Lower error sensitivity in random environment agrees with previous results in (Gonzalez Castro et al. 2014). Note that variability in movement outcomes can be both extrinsic (e.g., random condition in our experiment) and intrinsic (e.g., due to physiologic tremor). Our study focuses on the effect of the former. The effect of intrinsic motor variability on learning rate is unclear (He et al. 2016).

### Memory of Error Model

In stable condition, our results agree with the idea that error consistency sign modulates error sensitivity, offered as a part of “Memory of Error” model in (Herzfeld et al. 2014). However, the lack of error consistency dependence in the random condition (Fig 5) challenges the Memory of Error model (Albert et al. 2021; Herzfeld et al. 2014). A possible explanation for these differences is the presence of multiple targets and partial generalization in our study (Fig. 4). In contrast, the Memory of Error model does not account for target location and therefore assumes perfect generalization across targets, regardless of their spatial distance. Another possible explanation for these differences is the use of a joystick as controller in our study, which may potentially involve different adaptation mechanisms compared to the robotic arm used in (Herzfeld et al. 2014).

**Fig 5.**
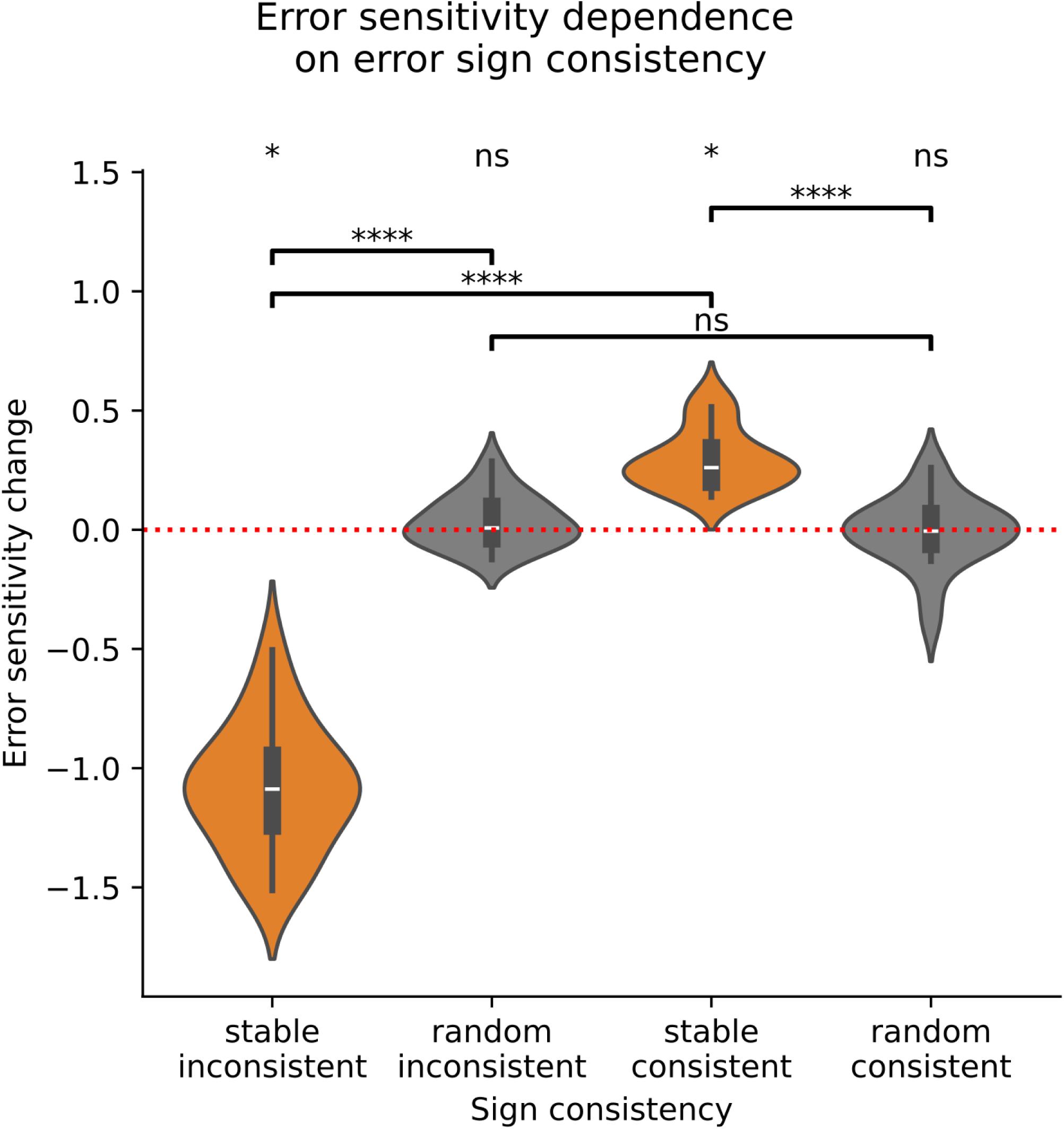
Changes of error sensitivity as a function of error sign consistency between consecutive trials, averaged within participant. Top line of significance corresponds to two-sided comparison against 0. All p-values are Holm-Bonferroni corrected. Note that even in stable perturbations, two consecutive errors can have inconsistent signs due to movement variability, especially close to the adaptation asymptote levels (see Fig. S5 for an individual participant error trace).

Note that in Fig 5, based on formula (2) we only check the validity of the *direction* of change of the consecutive error sensitivity values, not the magnitude. The formula (3) in (Herzfeld et al. 2014) prescribes the *magnitude* of change as well but we decided to omit comparing the magnitudes in our data with those prescribed by MoE model for two reasons 1) we judged that inconsistency in random environment observed in Fig 5 already demonstrates inconsistency of MoE model with our data 2) checking the magnitude values would require fitting the entirety of the generative MoE model to our data (which we tried and did not succeed probably due to the first reason).

### Error statistics and error sensitivity

As presented in Results section, we don’t find a universal relationship between the windowed error statistics and error sensitivity, in particular in random condition, unlike in (Tan et al. 2014). We describe several differences between that study and ours that can affect the comparison.

● Participants received online feedback of their movements and therefore could detect perturbation right on the very trial where it was introduced. We expect that it affects error sensitivity calculation, possibly in a way similar to the one presented in (Feulner et al. 2025).
● The distribution of random perturbations was different than in our study: there were only 7 different possibilities: 0, +/−12,+/−24,+/−40 degrees perturbations. This could affect the way perturbation variability is taken into account by the brain.
● The targets in Tan et al study were higher in number (eight) and they were disturbed differently than in our study: eight positions equally spaced around an invisible circle around the home position. It is possible that subjects credited some of the variance of their errors with the variance of the targets.
● Tan et al study featured 12 participants and it is generally known that motor adaptation rates and learning mechanisms involvement can vary between subjects (Moore and Cluff 2021).

### Implicit vs explicit learning

Motor adaptation involves two key processes: implicit learning and explicit strategy application. Implicit learning refers to the unconscious adjustment to a perturbation, aiming to partially compensate for it. Explicit strategy application, on the other hand, involves the conscious use of strategies such as re-aiming (Tsay et al. 2023).

Disentangling these two learning processes is challenging and requires careful experimental design (Tsay et al. 2023) Even with such efforts, the results can remain ambiguous (Maresch, Mudrik, and Donchin 2021). However, due to a large body of literature on the topic, some theoretical understanding starts to crystallize as well and certain learning phenomena are considered to be a feature of one process but not another.

Which mechanism is more prevalent in our setup is not exactly clear, but there are more reasons for it to be predominantly implicit, even though endpoint-only feedback often involves explicit learning (Tsay et al. 2023). The absence of savings (see next section) suggests lower involvement of explicit learning. One might argue that explicit learning has been partially disrupted by the random period. However, we observe evidence of spatial generalization during both the stable and random periods, and we find no differences in the shapes of the generalization curves. Given the unlikely role of explicit strategy in a random environment, the similarity in the shapes of the generalization curves leads us to hypothesize that implicit learning predominates in both the random and stable environments. Finally, a localized (unimodal, decaying to zero for large errors) spatial generalization profile (Fig S3, leftmost panel) is a feature of implicit learning (Tsay et al. 2023).

Note that spatial generalization in the context of environment stability was studied earlier (Fernandes, Stevenson, and Kording 2012) and was found to be similar in both high and low variance perturbation conditions. However the perturbation schedule there was different than in our experiment: first only one target was presented and then others. This way the possibility of forgetting what was learned for the training direction was not addressed. Besides that in (Avraham et al. 2020) it was already shown that error sensitivity is lower in low environmental consistency and the authors provided evidence that modulation was due to explicit learning. However, a more recent study (Albert et al. 2021) challenged this view showing that this effect is rather due to implicit learning and suggested that results in (Avraham et al. 2020) could be attributed to the relatively long (600 ms+) delay between the movement termination and endpoint feedback presentation.

Consistent with (Albert et al. 2021), we find that lower error consistency in the random environment does not reduce error sensitivity but rather suppresses its growth (Fig 2C, Fig 3). Dependence of error sensitivity on previous error has a familiar shape (Fig S3, first column) reported previously in (Herzfeld et al. 2014). We have not found a statistically significant difference between generalization curves in random and stable environments (Fig S3, first column).

### Savings

In our experiment the participants perceived the same perturbation twice: the first and last stable perturbation values are equal and also the second and third constant perturbation values are equal. In such situations, a phenomenon called “savings” may be observed (Smith et al. 2006): when re-experiencing a perturbation, adaptation could be faster. We did not find savings in our experiment. Error sensitivities were not statistically different between the first and last perturbation (two sample t-test, t=-1.90, p=1.47e-01, Bonferroni corrected) nor between the second and the third perturbation (t=1.33, p=4.01e-01, Bonferroni corrected).

Although still debated (Tsay et al. 2023) there is a hypothesis that savings occur exclusively in explicit strategy learning. In our data, however, no evidence of savings was observed, despite being expected. Error sensitivity values during the first and last perturbations were not significantly different (t = –1.90, p = 0.0733), nor were those during the second and third perturbations (t = 1.33, p = 0.200). This suggests that explicit learning was not the primary mechanism used by our participants.

There is evidence that savings can survive adaptation in the opposite direction (Malone et al. 2011; Sarwary, Selen, and Medendorp 2013). However, this evidence comes from the gait adaptation, where savings might be supported by different mechanisms. Moreover, the 2nd and 3rd perturbations in our experiment, which were not separated by an opposite consistent perturbation, still showed no evidence of savings in our analysis.

Savings suppression after a long washout of a special kind was observed earlier in (Kitago et al. 2013; Kojima, Iwamoto, and Yoshida 2004). In particular, (Kitago et al. 2013) report that long series of error clamp trials can suppress savings, and in our experiment the random environment can be considered a sort of random clamp environment, just that clamp direction changes on every trial. In (Albert et al. 2021) authors suggest a mechanism for savings suppression: error sensitivity update weight decay in Memory of Error model. Potentially, it could apply for our experiment as well because although the distribution of errors during random periods equaled the distribution of errors during the precedent stable period, the error consistency pattern was destroyed by trial permutation. However, since the Memory of Error model does not fully align with our data (see corresponding section above), it is unlikely to fully explain the lack of savings.

In addition to the random perturbations, there were three pauses lasting approximately one minute each. However, their influence on savings is likely less significant than that of the random period, which lasted nearly 15 minutes. The average pause duration was 44.7 seconds (SD = 13.4 seconds).

Another important consideration is connected to the results of (Hadjiosif, Morehead, and Smith 2023). The authors of that paper showed that motor adaptation can be split into a temporally persistent and temporary volatile component, the latter being defined as the learning that decays quickly (within 15-25 seconds). Somewhat counterintuitively, it is the volatile component that they show to be responsible for savings (the persistent one is responsible for long term retention). In our experiment, given that the targets appear in a pseudorandom order, in most cases the same target does not appear twice in a row. This creates a delay of multiple trials between the same target trials with mean time difference = 14.9 seconds (mean trail distance = 4 trials). While this number is slightly lower than 15-25 seconds mentioned in (Hadjiosif et al. 2023; Maurice Smith 2024), since the volatile adaptation decay is exponential and we have a relatively high motor noise in our experiment, we expect that this effect contributes to the absence of savings. The decay pattern of temporal generalization we observe in Fig 2B,D supports this idea (recall that we computed the quantity (2) looking at the subset of trials for which the target at trial n-k was the same as during trial k).

We would like to underline that this study was not designed to study savings specifically and further behavioral and modeling research needed to shed more light on the absence of savings in our setup.

### Trial dependence of Error Sensitivity

Trial dependence of error sensitivity is a matter of debate. Classical learning rate in motor adaptation was assumed to be constant, in particular, independent of error magnitude and history (Jordan and Rumelhart 1992; Kawato et al. 1987). Such theories do not capture the full variability observed in the data and one of the approaches to mitigate it was the introduction of learning rate dependence on error history (Herzfeld et al. 2014; Tan et al. 2014). This is important to mention however that in (Heald, Lengyel, and Wolpert 2021) authors proposed COIN model that gave an alternative explanation for results of (Herzfeld et al. 2014), based on context sequence switches. Such switches do incorporate error sensitivity dependence on error history in certain sense, but in a different way as suggested by (Herzfeld et al. 2014; Tan et al. 2014).

In (Wang et al. 2023) authors targeted the question of necessity of trial dependence of error sensitivity specifically and offered a model alternative to Memory of Error that is simpler and is able to explain their data as well as (Albert et al. 2021) data. However, it is not consistent with our data because in our experiment, error distributions between stable and random environments are designed to coincide perfectly and yet error sensitivity distributions between them are very different.

Continuous monotonous increase of error sensitivity within a stable environment was reported earlier in Experiment 6 of (Albert et al. 2021) using a different experimental controller and implicit learning-only design (achieved by reducing reacting time).

Beyond motor adaptation in non-motor cognitive science adaptive learning rates are also a topic of active research (Simoens et al. 2025).

### Generalization

A notable study on generalization was performed in (Donchin et al. 2003). There the generalization function shape was also obtained (cf Fig. 4) in both stable and random environments. Donchin study differs from our in several important ways:

● First it was done using a different experimental apparatus: force field robot. Unlike visuomotor adaptation which relies primarily on visual feedback, force field relies rather on proprioceptive feedback (Fleury, Prablanc, and Priot 2019; Pipereit, Bock, and Vercher 2006; Thomas and Bock 2012; Tzvi, Loens, and Donchin 2022; Zhou et al. 2024). Motor adaptation mechanisms with these two feedback types use different internal models and are thus not directly comparable.
● Second, since proprioceptive feedback is much faster than the visual one, participants could detect (and to certain extent identify) the perturbation on a given trial already on the same trial this perturbation was first applied. In our protocol with endpoint-only feedback the perturbation could only be perceived after the movement is made and thus its effect could only appear on the next trial.
● Third, the approach in (Donchin et al. 2003) assumes that error sensitivity depends exclusively on the limb velocity projection to so-called “basis elements” (a fixed number of functions that don’t change over the experiment). See eq (3) in (Donchin et al. 2003). Estimating the generalization function shape required making an assumption on the data generating mechanism and fitting a generative model (a system of difference equations). Our approach in contrast does not require any model fitting and allows for the error sensitivity to evolve freely without any assumptions.

Overall our research corroborates and complements findings in (Donchin et al. 2003).

### Controller type

The comparison of different motor adaptation studies should always be exercised with care, as various factors can influence the results. These include the type of controller used to guide the cursor (e.g., manipulandum, mouse, joystick, tablet, hand tracking), the type of perturbations (e.g., visuomotor rotations, visuomotor mirror reversal, visuomotor scaling, force field), the perturbation schedule (e.g., gradual, abrupt, random every N-th trial), the feedback presentation (e.g., online, online with delay, endpoint-only), and task timing (e.g., the presence of motor preparation and limits on the speed of the movement). Compared to manipulandums and hand tracking, joysticks are less commonly used as controllers in research.

Joysticks are routinely used in cognitive science research (Szul et al. 2020) and recently there has been a growing interest in using joysticks in motor adaptation experiments, because, not requiring strong activation of neck muscles, they simplify simultaneous neural recordings (Bracco et al. 2024; Reuter, Booms, and Leow 2022; Tan et al. 2014). Our study both allows us to compare how other generalization and error sensitivity studies with other controllers (Albert et al. 2021; Donchin et al. 2003; Herzfeld et al. 2014) generalize to our setup and also paves the way for future joint behavioral-neural analyses.

Moreover, spatial generalization patterns seem to strongly depend on the controller type, probably due to different relationships between proprioception-informed error and visual error and active vs passive behavior of the apparatus itself (Fleury et al. 2019). However the amount of decrease of error sensitivity with increasing target distance can strongly depend on the training type, in particular whether training was focused (involved similar movements) and with visual feedback or not (see Discussion in (Berniker et al. 2014)).

### Future directions

Overall, our results describe the error-sensitivity dynamic and its generalization in different environment, and partially overlap with existing theoretical approaches (Herzfeld et al. 2014; Tan et al. 2014). We expect that adding neural data to the analysis will allow us to explain more of the variance of error sensitivity values and will thus allow a better understanding of motor adaptation mechanisms, akin to (Bracco et al. 2024; Reuter et al. 2022). Magnetoencephalography data was collected simultaneously with the behavioral data analyzed in this study, and its analysis will be presented in future publications.

Recently in (Sugiyama, Schweighofer, and Izawa 2023) a new model of motor adaptation was suggested that links adaptation rate to reward. There the reward was given explicitly to modulate learning. It would be interesting to explore whether this approach could be adapted to situations without explicit rewards, where participants may rely on internal or self-generated reinforcement. However, modeling such internal reward signals remains a complex challenge (Shmuelof et al. 2012; Todorov et al. 2019).

## Acknowledgements

We thank D. Herzfeld, O. Abdoun and G. Marrelec for their valuable advice and the EDUWELL team at the Lyon Neuroscience Research Center (CRNL) for general support.

This research was supported in part by the Intramural Research Program of the National Institutes of Health (NIH). The contributions of the NIH authors were made as part of their official duties as NIH federal employees, are in compliance with agency policy requirements, and are considered Works of the United States Government. However, the findings and conclusions presented in this paper are those of the authors and do not necessarily reflect the views of the NIH or the U.S. Department of Health and Human Services.

## References

1. Albert, Scott T., Jihoon Jang, Hannah R. Sheahan, Lonneke Teunissen, Koenraad Vandevoorde, David J. Herzfeld, and Reza Shadmehr. 2021. “An Implicit Memory of Errors Limits Human Sensorimotor Adaptation.” Nature Human Behaviour 5(7):920–34. doi:10.1038/s41562-020-01036-x.

2. Avraham, Guy, Matan Keizman, and Lior Shmuelof. 2020. “Environmental Consistency Modulation of Error Sensitivity during Motor Adaptation Is Explicitly Controlled.” Journal of Neurophysiology 123(1):57–69. doi:10.1152/jn.00080.2019.

3. Berniker, Max, David W. Franklin, J. Randall Flanagan, Daniel M. Wolpert, and Konrad Kording. 2014. “Motor Learning of Novel Dynamics Is Not Represented in a Single Global Coordinate System: Evaluation of Mixed Coordinate Representations and Local Learning.” Journal of Neurophysiology 111(6):1165–82. doi:10.1152/jn.00493.2013.

4. Bracco, Martina, Varsha Vasudevan, Vridhi Rohira, Quentin Welniarz, Mihoby Razafinimanana, Alienor Richard, Christophe Gitton, Sabine Meunier, Antoni Valero-Cabré, Denis Schwartz, Traian Popa, and Cécile Gallea. 2024. “Emergence of Pre-Movement Beta Activity with Stable Sensorimotor Predictions to Facilitate Motor Adjustments.” 2024.12.23.630078.

5. Brayanov, Jordan B., Daniel Z. Press, and Maurice A. Smith. 2012. “Motor Memory Is Encoded as a Gain-Field Combination of Intrinsic and Extrinsic Action Representations.” Journal of Neuroscience 32(43):14951–65. doi:10.1523/JNEUROSCI.1928-12.2012.

6. Donchin, Opher, Joseph T. Francis, and Reza Shadmehr. 2003. “Quantifying Generalization from Trial-by-Trial Behavior of Adaptive Systems That Learn with Basis Functions: Theory and Experiments in Human Motor Control.” Journal of Neuroscience 23(27):9032–45. doi:10.1523/JNEUROSCI.23-27-09032.2003.

7. Fernandes, Hugo L., Ian H. Stevenson, and Konrad P. Kording. 2012. “Generalization of Stochastic Visuomotor Rotations.” PloS One 7(8):e43016. doi:10.1371/journal.pone.0043016.

8. Feulner, Barbara, Matthew G. Perich, Lee E. Miller, Claudia Clopath, and Juan A. Gallego. 2025. “A Neural Implementation Model of Feedback-Based Motor Learning.” Nature Communications 16(1):1805. doi:10.1038/s41467-024-54738-5.

9. Fleury, Lisa, Claude Prablanc, and Anne-Emmanuelle Priot. 2019. “Do Prism and Other Adaptation Paradigms Really Measure the Same Processes?” Cortex 119:480–96. doi:10.1016/j.cortex.2019.07.012.

10. Gonzalez Castro, Luis Nicolas, Alkis M. Hadjiosif, Matthew A. Hemphill, and Maurice A. Smith. 2014. “Environmental Consistency Determines the Rate of Motor Adaptation.” Current Biology 24(10):1050–61. doi:10/f54fbf.

11. Hadjiosif, Alkis M., J. Ryan Morehead, and Maurice A. Smith. 2023. “A Double Dissociation between Savings and Long-Term Memory in Motor Learning.” PLOS Biology 21(4):e3001799. doi:10.1371/journal.pbio.3001799.

12. He, Kang, You Liang, Farnaz Abdollahi, Moria Fisher Bittmann, Konrad Kording, and Kunlin Wei. 2016. “The Statistical Determinants of the Speed of Motor Learning.” PLOS Computational Biology 12(9):e1005023. doi:10.1371/journal.pcbi.1005023.

13. Heald, James B., Máté Lengyel, and Daniel M. Wolpert. 2021. “Contextual Inference Underlies the Learning of Sensorimotor Repertoires.” Nature 600(7889):489–93. doi:10.1038/s41586-021-04129-3.

14. Herzfeld, David J., Pavan A. Vaswani, Mollie K. Marko, and Reza Shadmehr. 2014. “A Memory of Errors in Sensorimotor Learning.” Science 345(6202):1349–53. doi:10.1126/science.1253138.

15. Jordan, Michael I., and David E. Rumelhart. 1992. “Forward Models: Supervised Learning with a Distal Teacher.” Cognitive Science 16(3):307–54. doi:10.1016/0364-0213(92)90036-T.

16. Jordan, Michael I., and David E. Rumelhart. 1995. “Forward Models: Supervised Learning with a Distal Teacher.” in Backpropagation. Psychology Press.

17. Kawato, M., Kazunori Furukawa, and R. Suzuki. 1987. “A Hierarchical Neural-Network Model for Control and Learning of Voluntary Movement.” Biological Cybernetics 57(3):169–85. doi:10.1007/BF00364149.

18. Kitago, Tomoko, Sophia Ryan, Pietro Mazzoni, John Krakauer, and Adrian Haith. 2013. “Unlearning versus Savings in Visuomotor Adaptation: Comparing Effects of Washout, Passage of Time, and Removal of Errors on Motor Memory.” Frontiers in Human Neuroscience 7. https://www.frontiersin.org/articles/10.3389/fnhum.2013.00307.

19. Kojima, Yoshiko, Yoshiki Iwamoto, and Kaoru Yoshida. 2004. “Memory of Learning Facilitates Saccadic Adaptation in the Monkey.” Journal of Neuroscience 24(34):7531–39. doi:10.1523/JNEUROSCI.1741-04.2004.

20. Krakauer, John W. 2006. “Motor Learning: Its Relevance to Stroke Recovery and Neurorehabilitation.” Current Opinion in Neurology 19(1):84–90. doi:10.1097/01.wco.0000200544.29915.cc.

21. Krakauer, John W., and S. Thomas Carmichael. 2022. Broken Movement: The Neurobiology of Motor Recovery after Stroke. MIT Press.

22. Krakauer, John W., Pietro Mazzoni, Ali Ghazizadeh, Roshni Ravindran, and Reza Shadmehr. 2006. “Generalization of Motor Learning Depends on the History of Prior Action.” PLOS Biology 4(10):e316. doi:10.1371/journal.pbio.0040316.

23. Malone, Laura A., Erin V. L. Vasudevan, and Amy J. Bastian. 2011. “Motor Adaptation Training for Faster Relearning.” Journal of Neuroscience 31(42):15136–43. doi:10.1523/JNEUROSCI.1367-11.2011.

24. Maresch, Jana, Liad Mudrik, and Opher Donchin. 2021. “Measures of Explicit and Implicit in Motor Learning: What We Know and What We Don’t.” Neuroscience & Biobehavioral Reviews 128:558–68. doi:10.1016/j.neubiorev.2021.06.037.

25. Maurice Smith. 2024. “Maurice Smith: Cerebellum and Human Motor Memory.” Presented at the Shadmehr Lab Seminar series (available on YouTube), December 10, online.

26. Mazzoni, Pietro, and John W. Krakauer. 2006. “An Implicit Plan Overrides an Explicit Strategy during Visuomotor Adaptation.” Journal of Neuroscience 26(14):3642–45. doi:10.1523/JNEUROSCI.5317-05.2006.

27. Moore, Robert T., and Tyler Cluff. 2021. “Individual Differences in Sensorimotor Adaptation Are Conserved Over Time and Across Force-Field Tasks.” Frontiers in Human Neuroscience 15:692181. doi:10.3389/fnhum.2021.692181.

28. Morehead, J. Ryan, and Jean-Jacques Orban de Xivry. 2021. *A Synthesis of the Many Errors and Learning Processes of Visuomotor Adaptation*. *preprint*. Neuroscience. doi:10.1101/2021.03.14.435278.

29. Pipereit, Katja, Otmar Bock, and Jean-Louis Vercher. 2006. “The Contribution of Proprioceptive Feedback to Sensorimotor Adaptation.” Experimental Brain Research 174(1):45–52. doi:10.1007/s00221-006-0417-7.

30. Reuter, Eva-Maria, Arthur Booms, and Li-Ann Leow. 2022. “Using EEG to Study Sensorimotor Adaptation.” Neuroscience & Biobehavioral Reviews 134:104520. doi:10.1016/j.neubiorev.2021.104520.

31. Sarwary, A. M. E., L. P. J. Selen, and W. P. Medendorp. 2013. “Vestibular Benefits to Task Savings in Motor Adaptation.” Journal of Neurophysiology 110(6):1269–77. doi:10.1152/jn.00914.2012.

32. Shmuelof, Lior, Vincent S. Huang, Adrian M. Haith, Raymond J. Delnicki, Pietro Mazzoni, and John W. Krakauer. 2012. “Overcoming Motor ‘Forgetting’ through Reinforcement of Learned Actions.” Journal of Neuroscience 32(42):14617–21. doi:10.1523/jneurosci.2184-12.2012.

33. Simoens, Jonas, Senne Braem, Pieter Verbeke, Haopeng Chen, Stefania Mattioni, Mengqiao Chai, Nicolas W. Schuck, and Tom Verguts. 2025. “Two Time Scales of Adaptation in Human Learning Rates.” 2025.06.05.658048.

34. Smith, Maurice A., Ali Ghazizadeh, and Reza Shadmehr. 2006. “Interacting Adaptive Processes with Different Timescales Underlie Short-Term Motor Learning” edited by J. Ashe. PLoS Biology 4(6):e179. doi:10.1371/journal.pbio.0040179.

35. Sugiyama, Taisei, Nicolas Schweighofer, and Jun Izawa. 2023. “Reinforcement Learning Establishes a Minimal Metacognitive Process to Monitor and Control Motor Learning Performance.” Nature Communications 14(1):3988. doi:10.1038/s41467-023-39536-9.

36. Szul, Maciej J., Aline Bompas, Petroc Sumner, and Jiaxiang Zhang. 2020. “The Validity and Consistency of Continuous Joystick Response in Perceptual Decision-Making.” Behavior Research Methods 52(2):681–93. doi:10.3758/s13428-019-01269-3.

37. Tan, Huiling, Ned Jenkinson, and Peter Brown. 2014. “Dynamic Neural Correlates of Motor Error Monitoring and Adaptation during Trial-to-Trial Learning.” Journal of Neuroscience 34(16):5678–88. doi:10.1523/JNEUROSCI.4739-13.2014.

38. Tanaka, Hirokazu, and Terrence J. Sejnowski. 2014. “Motor Adaptation and Generalization of Reaching Movements Using Motor Primitives Based on Spatial Coordinates.” Journal of Neurophysiology 113(4):1217–33. doi:10/f62679.

39. Thomas, Monika, and Otmar Bock. 2012. “Concurrent Adaptation to Four Different Visual Rotations.” Experimental Brain Research 221(1):85–91. doi:10.1007/s00221-012-3150-4.

40. Thoroughman, Kurt A., and Reza Shadmehr. 2000. “Learning of Action through Adaptive Combination of Motor Primitives.” Nature 407(6805):742–47. doi:10.1038/35037588.

41. Todorov, Dmitrii. 2025. “Motor_Adaptation_VMR_Joystick_Stable_and_Random.” OSF. doi:10.17605/OSF.IO/6J8SM.

42. Todorov, Dmitrii I., Robert A. Capps, William H. Barnett, Elizaveta M. Latash, Taegyo Kim, Khaldoun C. Hamade, Sergey N. Markin, Ilya A. Rybak, and Yaroslav I. Molkov. 2019. “The Interplay between Cerebellum and Basal Ganglia in Motor Adaptation: A Modeling Study.” Plos One 14(4):e0214926.

43. Tsay, Jonathan, Richard Ivry, John W. Krakauer, Adrian M. Haith, Guy Avraham, and Hyosub E. Kim. 2023. “Strategy Processes in Sensorimotor Learning: Reasoning, Refinement, and Retrieval.”

44. Tzvi, Elinor, Sebastian Loens, and Opher Donchin. 2022. “Mini-Review: The Role of the Cerebellum in Visuomotor Adaptation.” The Cerebellum 21(2):306–13. doi:10.1007/s12311-021-01281-4.

45. Vallat, Raphael. 2018. “Pingouin: Statistics in Python.” Journal of Open Source Software 3(31):1026. doi:10.21105/joss.01026.

46. Wang, Tianhe, Guy Avraham, Jonathan S. Tsay, Sabrina J. Abram, and Richard B. Ivry. 2023. “Perturbation Variability Does Not Influence Implicit Sensorimotor Adaptation.” 2023.01.27.525949.

47. Zhou, Weiwei, Emma Monsen, Kareelynn Donjuan Fernandez, Katelyn Haly, Elizabeth A. Kruse, and Wilsaan M. Joiner. 2024. “Motion State-Dependent Motor Learning Based on Explicit Visual Feedback Has Limited Spatiotemporal Properties Compared with Adaptation to Physical Perturbations.” Journal of Neurophysiology 131(2):278–93. doi:10.1152/jn.00198.2023.

