## Appendix for "Modulation of error sensitivity during motor learning across time, space and environment variability"

#### Additional figures

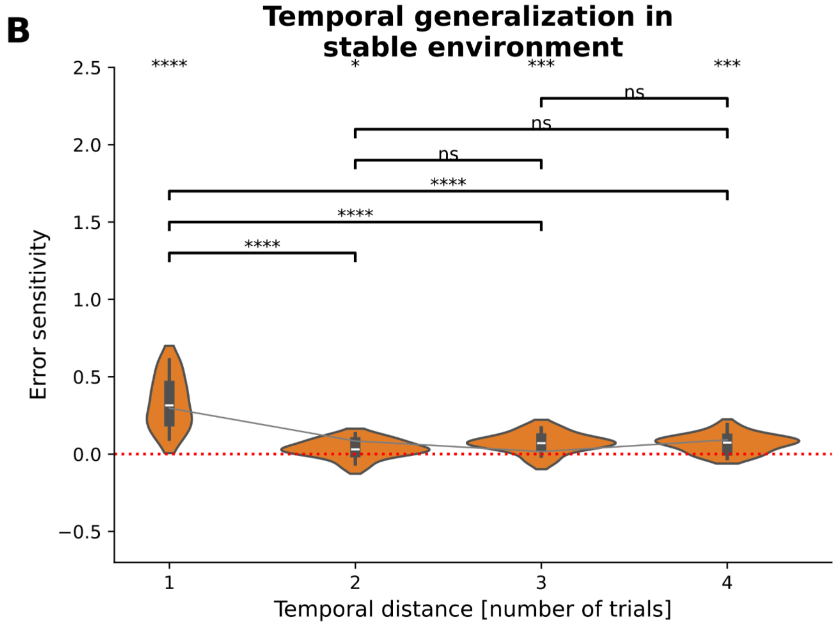


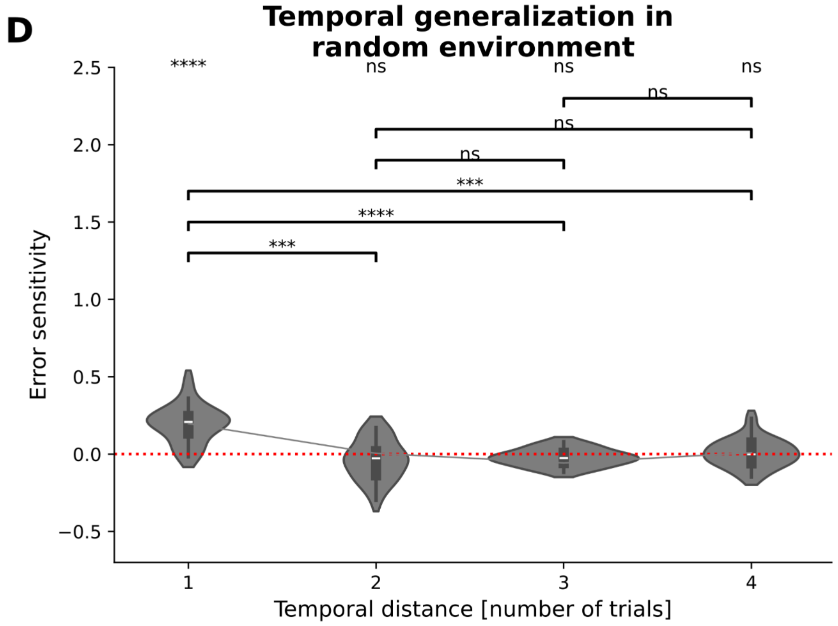

Fig S1: Temporal generalization pattern for ES_k computed using all trials, including those with different targets.

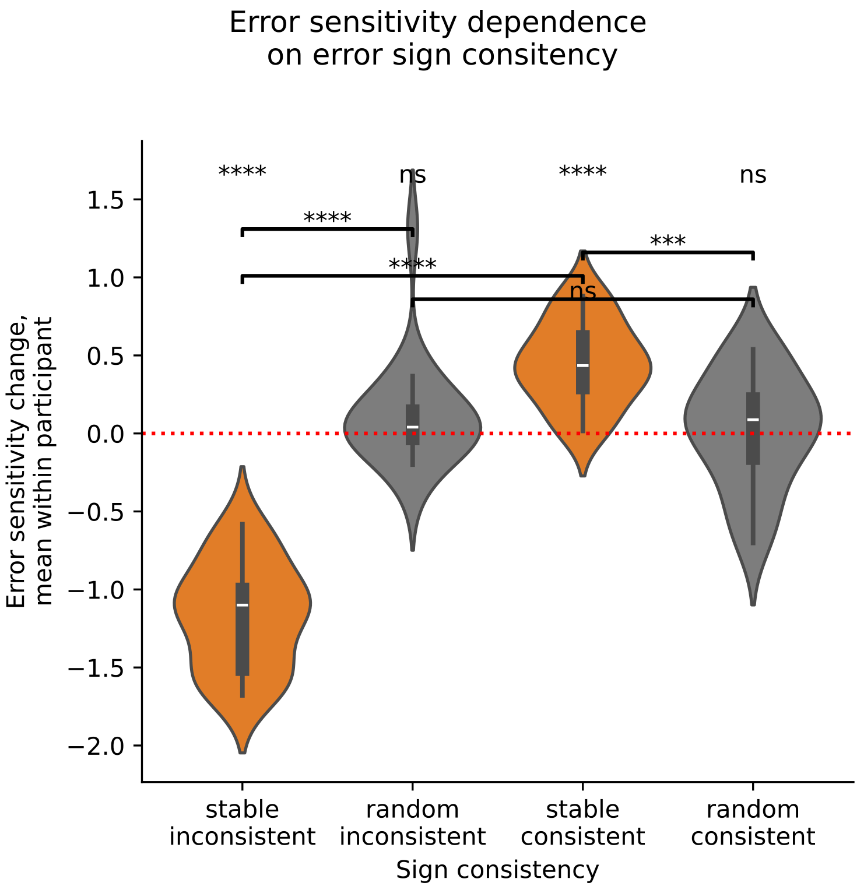


Fig S2: Changes of error sensitivity as a function of error sign consistency between consecutive trials, averaged within participant. Top line of significance codes corresponds to two-sided comparison against 0. Allo p-values are Holm-Bonferroni corrected. Analysis was done excluding trials whether either previous or pre-previous errors were smaller than the standard deviation of the error in the no-perturbation stage.

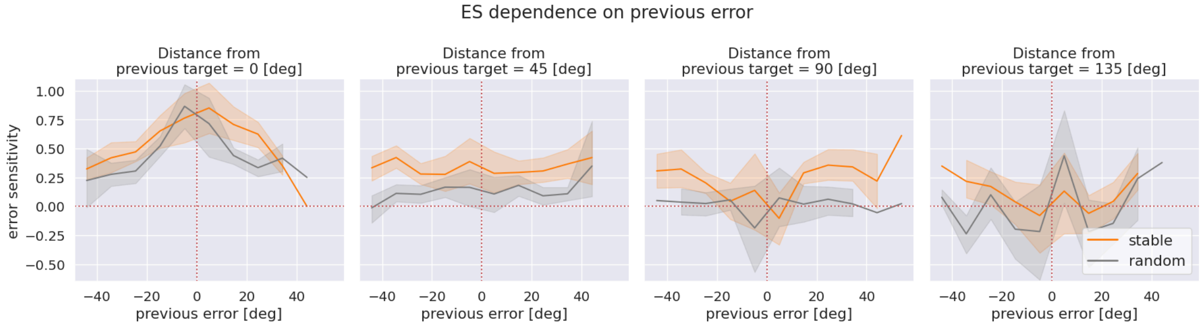


Fig S3. Dependence of error sensitivity on previous error, averages within subject. Bins with less than three datapoints were excluded. Previous error was binned into 15 bins.

### Error sensitivity within-target

One can compute error sensitivity using modified formula (1): instead of simply taking consecutive trials one can only consider trials on which the same target was presented. When computed this way, error sensitivity evolves as shown in Fig S4, qualitatively similar to the values computed used in the main text (see Fig 2B). We tried to perform our analysis on these error sensitivity values as well and we got results qualitatively similar to those described in the main manuscript.

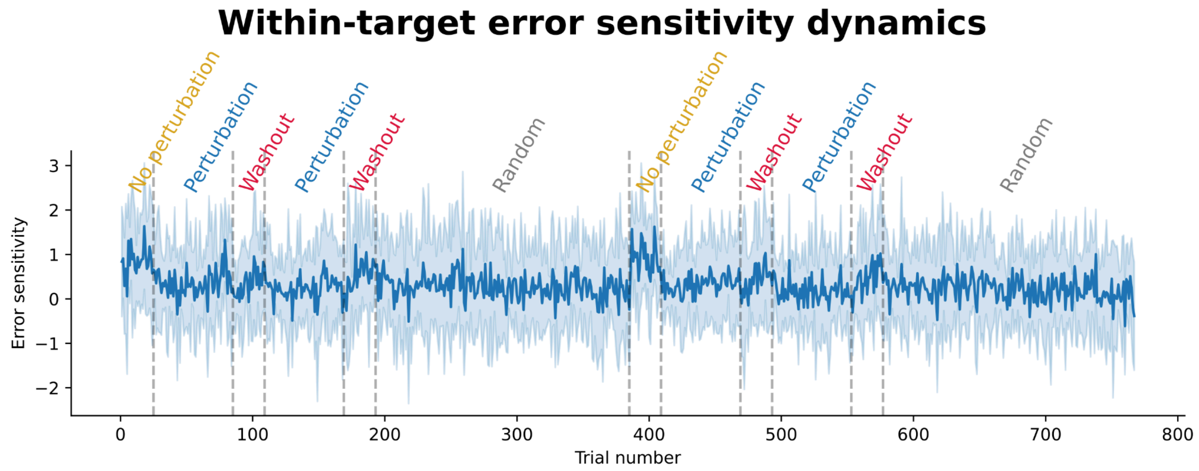


Figure S4. Evolution of within-target error sensitivity across trials (average across subjects, shading shows standard deviation).

#### Additional experiment stage statistics

We noticed that a spearman slope for the third perturbation (see Fig 2A) was significantly higher than for the fourth perturbation (t=4.59, p=4e-4, Bonferroni corrected). Since the two perturbations in question are the opposite to each other, we did not include this result in the discussion of savings.

We also report comparison of sub-stages (we separate the experiment not in 2 or 4 like in the main text, but in 12 stages). We found that error sensitivity in first and second random appearances were not different (p-value = 0.94). Also last perturbation has lower error sensitivity compared to first and third perturbations (p-values 0.035 and 0.016. And the first washout has lower error sensitivity compared to the second one (p-value 0.014).

We have also repeated part of the analysis (Savings test and Fig 2) when error sensitivity was calculated within-target, i.e. assuming targets involve completely separate memories, i.e. that in formula 1 the previous trial is actually the previous trial *when the same target was observed*. The results were qualitatively the same. We have also repeated the analysis with the retention factor in formula 1 equal to 1, again with the same results.

### Tan-like Windowed Error Statistics

One can the formula (4) in the manuscript in the following way: consider the history of ***N*** last trials and the following formula:

$$AR_{N,m,s}(n) = \frac{\mu_N^m(n)}{\sigma_N^s(n)}, \tag{S1}$$

Where n is the trial number, *N* takes values from 3 to 39, the exponents *m* and *s* are integers taking values from [0,1,2]. This way formula S1 allows to describe the following statistics:

- the standard deviation of previous errors,
- the square of the mean of previous errors,
- the inverse standard deviation of previous errors,
- the ratio of mean to standard deviation,
- the ratio of mean to variance and
- the ratio of mean squared to variance (Formula 4).

We performed the same analysis as in “Error sensitivity relation to statistics of recent errors” section of results and reported the summary in Table S1. It is easy to see that no statistic is significantly positively correlated in both environments. We put the full table (1776 entries) in .csv format with p-values and mean R2 values in the github repository github.com. See also Fig. S6 and S7.

| | window statistic formula | history length intervals | signed | environment |
|---|---|---|---|---|
| 0 | 1/std | 3-39 | False | stable |
| 1 | 1/std | 4 | True | random |
| 2 | 1/std | 6-39 | True | stable |
| 3 | mean/var | 4-39 | False | stable |
| 4 | mean^2 | 14-15 | True | random |
| 5 | mean^2/var | 12, 14 | True | random |
| 6 | std | 13-15, 17-20 | False | random |
| 7 | std | 19 | True | random |
| 8 | std | 3 | True | stable |

Table S1: Summary for correlation analysis of relation between windowed error statistics AR(N) and error sensitivity. We performed the correlations for various windowed statistics (formula S1) for various history lengths between 3 and 39 (second column, intervals are inclusive), for signed and absolute values of error (third column) for random and stable environments separately (fourth column). Only the statistics whose Spearman rhos computed within participants were significantly positive are shown. Abbreviations used: std = standard deviation, var = variance.

#### Single Subject Error Sensitivity

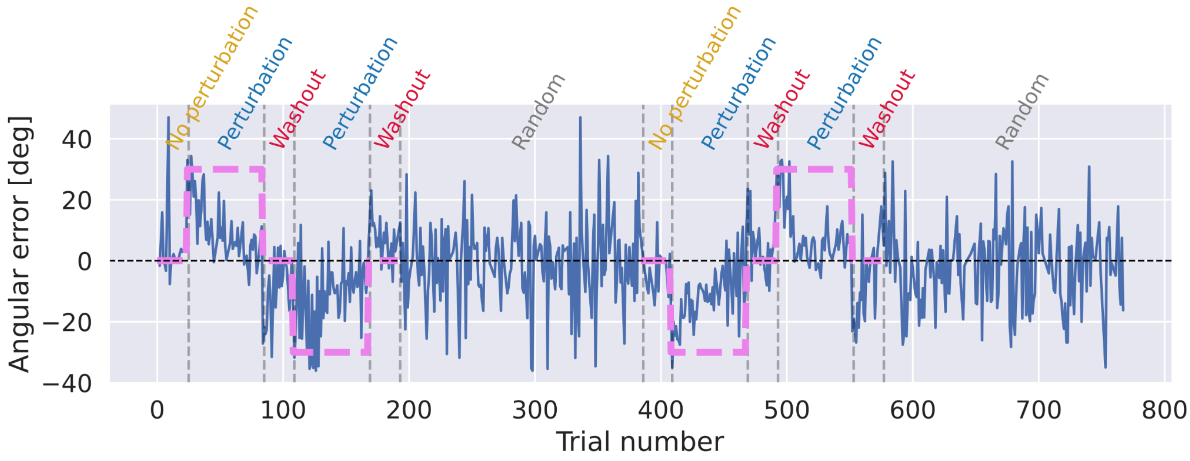


**Figure S5**. Error dynamics for a single participant. Purple line represents perturbation, text on top of A denotes experimental conditions.

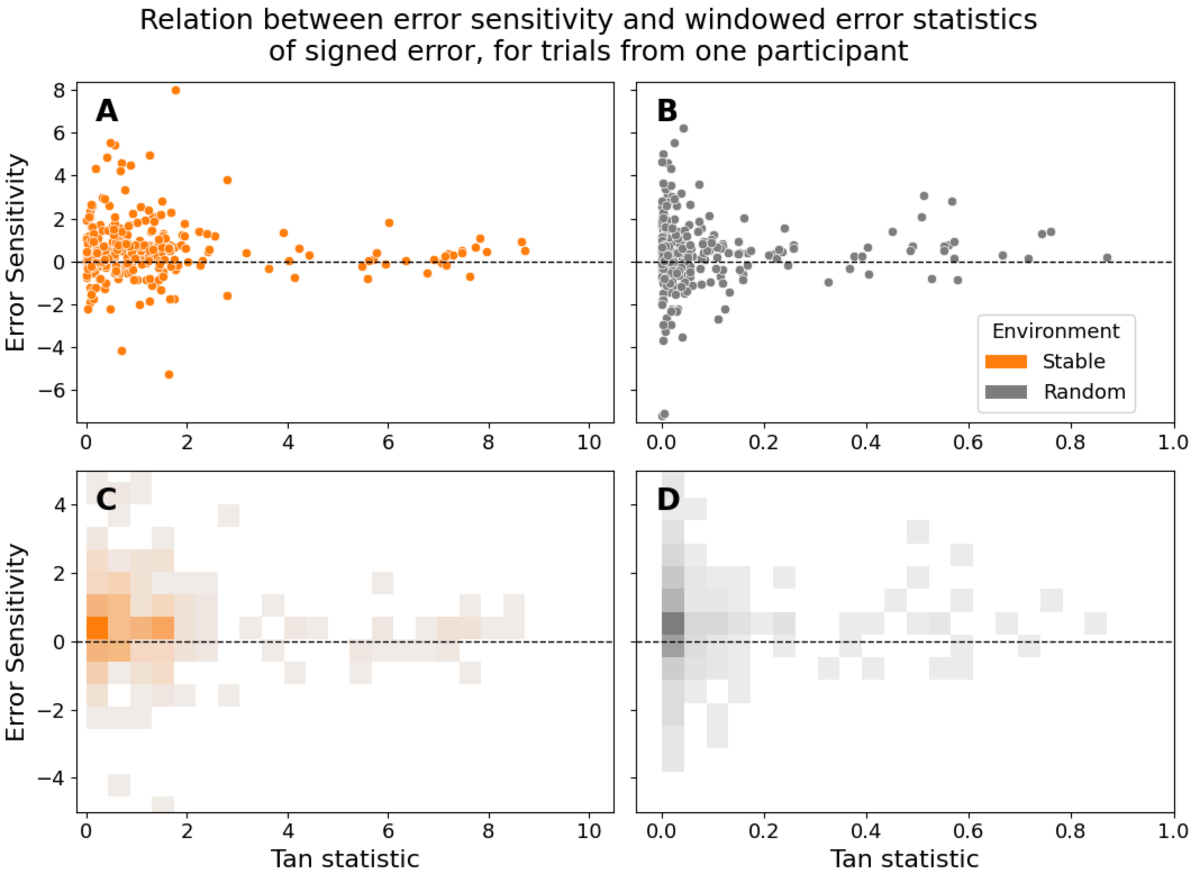


**Figure S6**. Relation between error sensitivity and Tan statistic (formula (4) in the manuscript), for one representative participant. **A,B** scatter plots in stable and random environments **C,D** histogram in stable and random environments.

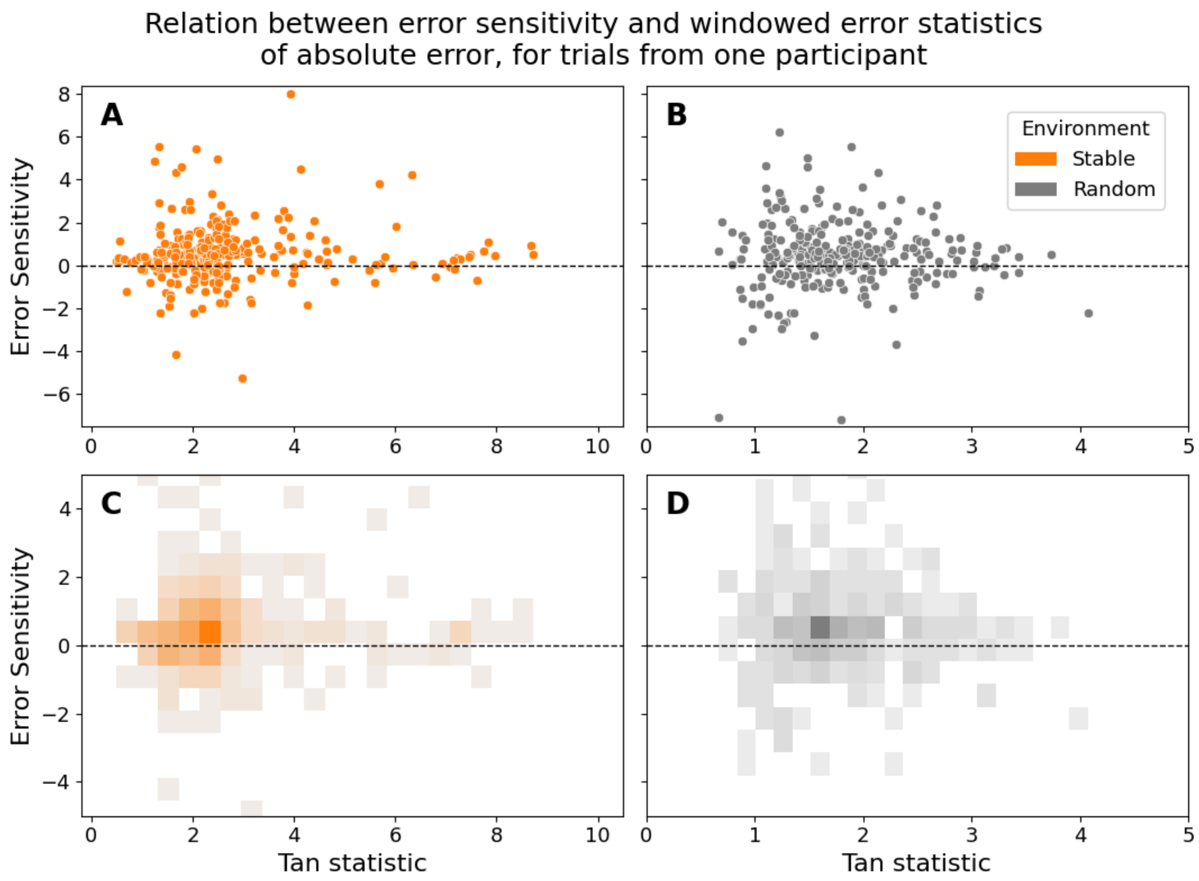


**Figure S7**. Relationship between error sensitivity and Tan statistic (formula (4) in the manuscript with absolute error values used for the mean and standard deviation computation), for one representative participant (same as in Fig. S6). **A,B** scatter plots in stable and random environments **C,D** two-dimensional histogram in stable and random environments.

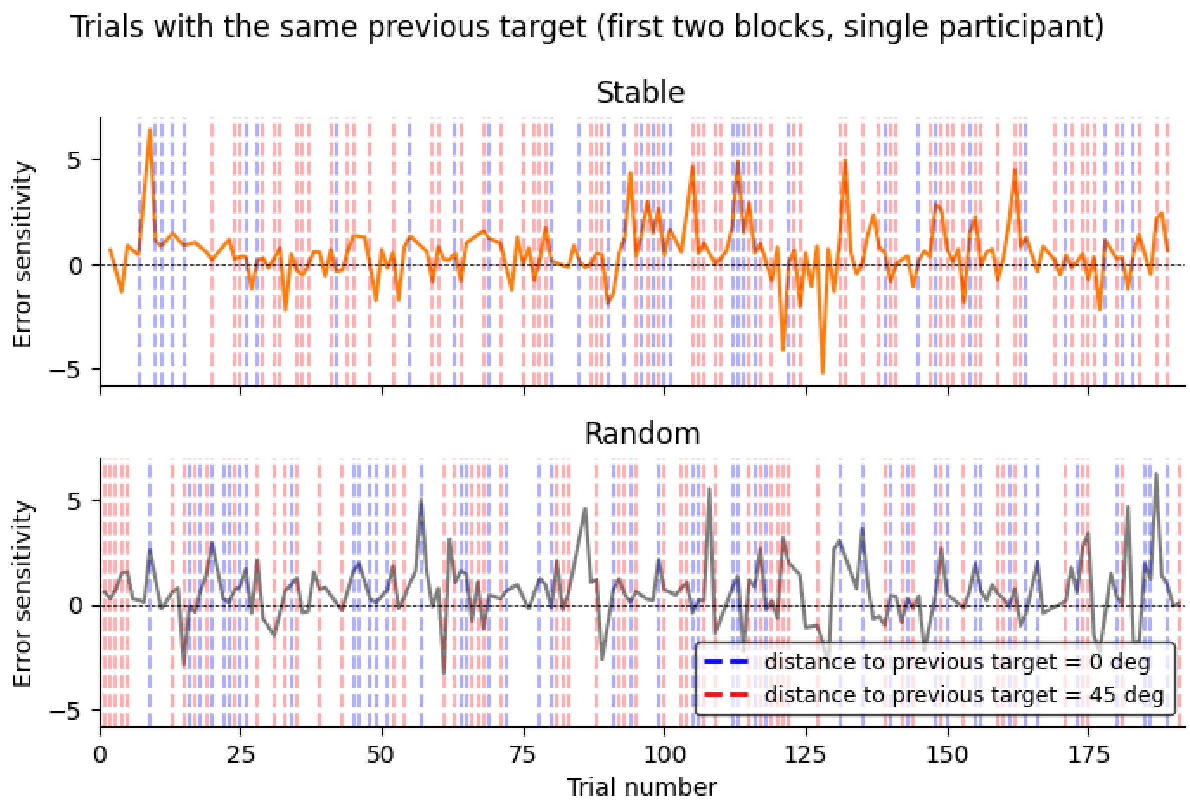


**Figure S8.** Temporal location of trials with previous target being at a given distance

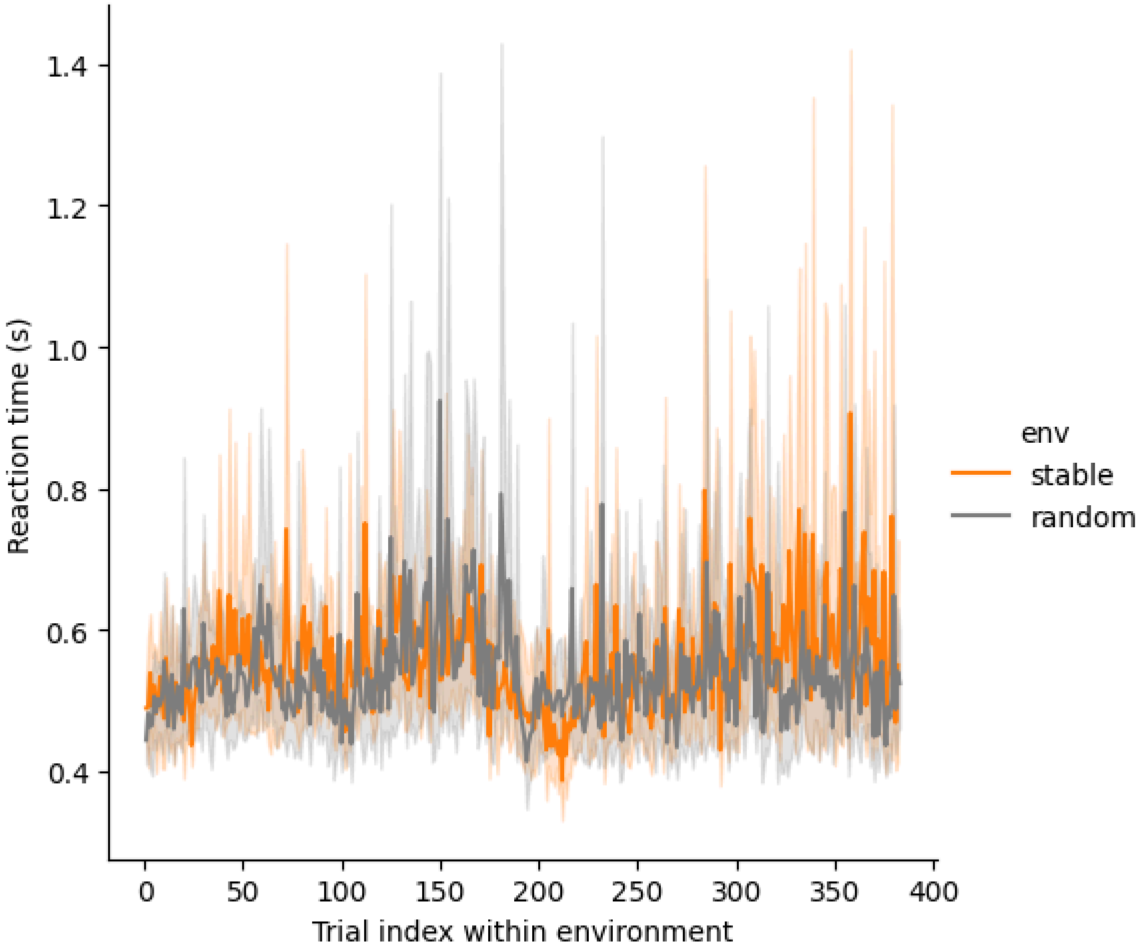


**Figure S9**. Reaction time evolution within each stability environment (both environment sub-blocks concatenated). Mean of average reaction time within subject = 0.54s +/- 0.12s.

| Source | SS | DF | MS | F | p-unc | p-GG-corr | ng2 | eps | sphericity | W-spher | p-spher |
|---|---|---|---|---|---|---|---|---|---|---|---|
| ps2_ | 8.527211 | 3 | 2.842404 | 85.695098 | 4.171004e-21 | 1.098059e-14 | 0.74884 | 0.660176 | False | 0.347063 | 0.002173 |
| Error | 1.890622 | 57 | 0.033169 | NaN | NaN | NaN | NaN | NaN | NaN | NaN | NaN |

**Table S2**. results one way repeated measures ANOVA comparing effect of sub-condition (ps2_) on error sensitivity

| Source | SS | ddof1 | ddof2 | MS | F | p-unc | p-GG-corr | ng2 | eps |
|---|---|---|---|---|---|---|---|---|---|
| env | 0.715429 | 1 | 19 | 0.715429 | 30.262084 | 2.628694e-05 | 2.628694e-05 | 0.100086 | 1.000000 |
| dist_rad_from_prevtgt_shiftrespect | 8.208524 | 3 | 57 | 2.736175 | 71.591798 | 2.533952e-19 | 6.407658e-15 | 0.560645 | 0.742735 |
| env * dist_rad_from_prevtgt_shiftrespect | 0.028399 | 3 | 57 | 0.009466 | 0.419325 | 7.398195e-01 | 6.860869e-01 | 0.004395 | 0.759186 |

**Table S3**. results two way repeated measures ANOVA comparing effect of condition (env) and previous target distance (dist_rad_from_prevtgt_shiftrespect) on error sensitivity
